# Hepatic stellate cells maintain liver homeostasis through paracrine neurotrophin-3 signaling

**DOI:** 10.1101/2023.03.03.531042

**Authors:** Vincent Quoc-Huy Trinh, Ting-Fang Lee, Sara Lemoinne, Kevin C. Ray, Maria D. Ybanez, Takuma Tsuchida, James K. Carter, Judith Agudo, Brian D. Brown, Kemal M. Akat, Scott L. Friedman, Youngmin A. Lee

**Affiliations:** Department of Surgery, Vanderbilt University Medical Center; Nashville, TN, USA; Division of Liver Diseases, Icahn School of Medicine at Mount Sinai; New York, NY, USA; Cancer Immunology and Virology, Dana-Farber Cancer Institute, Harvard Medical School; Boston, MA, USA; Icahn Genomics Institute, Icahn School of Medicine at Mount Sinai, New York, NY, USA; Precision Immunology Institute, Icahn School of Medicine at Mount Sinai, New York, NY, USA; Division of Cardiology, Department of Medicine, Vanderbilt University Medical Center; Nashville, TN, USA

## Abstract

Organ homeostasis is maintained by regulated proliferation of distinct cell populations. In mouse liver, cyclin D1-positive hepatocytes in the midlobular zone repopulate the parenchyma at a constant rate to preserve liver homeostasis. The mitogenic cues that underlie this process are unknown. Hepatic stellate cells, the liver’s pericytes, are in close proximity to hepatocytes and have been implicated in supporting hepatocyte proliferation, but their role in liver homeostasis is unknown. Here, we employ a T cell-mediated hepatic stellate cell ablation model to remove nearly all hepatic stellate cells in the murine liver, enabling the unbiased characterization of hepatic stellate cell functions. In the normal murine liver, complete loss of hepatic stellate cells persists for up to 6 weeks and reduces liver mass. Our results show that hepatic stellate cells induce cyclin D1 in midlobular hepatocytes by release of neurotrophin-3 to promote hepatocyte proliferation via tropomyosin receptor kinase B signaling. These findings establish that hepatic stellate cells form the niche for midlobular hepatocytes and reveal a novel hepatocyte growth factor signaling pathway.

**One-Sentence Summary:** Hepatic stellate cells provide mitogenic cues for midlobular hepatocyte proliferation and metabolic zonation by secreting neurotrophin-3.

## Main Text

Metabolic liver zonation is critical for liver function (*1*). The liver is organized into functional units, liver lobules, in which hepatocytes express metabolic genes in a positional manner. Traditionally, the numbering of liver zones follows the direction of the hepatic blood flow, from the periportal (zone 1) to the midlobular (zone 2), then to the pericentral (zone 3) hepatocytes. Spatial transcriptomics studies in healthy and injured livers suggest that hepatocytes are organized in up to 9 lobular layers with a gradient of metabolic specialization across these zones, reinforcing the importance of zonation to liver function (*2–5*).

In healthy liver, only a subset of hepatocytes proliferates at any given time to steadily replenish and maintain a defined organ size and hepatocellular mass. The cellular source of repopulating hepatocytes in this setting had been debated (*6–10*), but recent studies using sophisticated labeling approaches provided strong evidence that midlobular zone 2 hepatocytes proliferate to maintain liver homeostasis (*11, 12*). Central to this finding is cyclin D1 (CCND1), a cell cycle protein that promotes progression from G1 to S phase (*13*). CCND1 is constitutively nuclear and expressed in most midlobular hepatocytes, and a subset of CCND1 positive midlobular hepatocytes are proliferating at any given time. Hepatocyte-specific deletion of CCND1 leads to loss of proliferation in midlobular hepatocytes (*11*), and acute deletion of CCND1 delays liver regeneration (*14*), underscoring the pivotal role of CCND1-positive midlobular hepatocytes.

Although advanced models have provided more insight into the source of proliferating hepatocytes during homeostasis (*11, 12*), the cues determining this process and the mitogenic signals driving hepatocyte proliferation in zone 2 remain unknown (*15, 16*). Hepatic stellate cells (HSC) are the liver’s pericytes and reside in the space of Disse in direct contact with hepatocytes. HSCs release growth factors during development, including Wnt9a (*17*), hepatocyte growth factor (HGF) (*18*) and others (*19–22*). However, disruption of Wnt signaling in HSCs does not alter zonation, liver proliferation, or fibrosis (*23*). Similarly, deletion of HGF in HSCs did not abrogate tumorigenesis in a murine model of cholangiocarcinoma, a liver cancer arising from biliary epithelial cells (*24*). Because HSCs can differentiate into fibrogenic myofibroblasts during liver injury, much research has focused on their fibrogenic effects rather than on their role in liver homeostasis.

### Statement of specific scope of the study

The aim of this study was to develop a highly efficient HSC ablation model and to determine the contribution of HSCs to liver homeostasis.

## Results

### Near complete HSC depletion using eGFP-specific Jedi T cells

Agudo and colleagues developed eGFP-specific T cells that enable targeted cell depletion in eGFP-expressing mice (*25, 26*). *Just eGFP death-inducing* (Jedi) mice harbor T cells with T cell receptors (TCR) specific for the immunodominant epitope of eGFP presented by MHC class I (MHC-I). Adoptive transfer of Jedi T cells into an eGFP-reporter mouse with the same MHC-I haplotype H-2K^d^ efficiently ablates eGFP-positive cells within days. Single-cell RNA-seq (scRNA-seq) analyses from human and mouse livers have revealed the heterogeneity and spatial zonation of HSCs (*27–31*). Because analysis of published scRNA-seq datasets showed high expression and specificity of *platelet-derived growth factor receptor β* (*Pdgfrb*) in HSCs in both injured and healthy livers (**Suppl. Fig. 1A**), we used *Pdgfrb*-BAC-eGFP transgenic mice (henceforth called *Pdgfrb*-GFP), which express eGFP in HSCs (*28, 32*). Immunostaining for GFP in livers of *Pdgfrb*-GFP mice showed staining only in small non-parenchymal cells (**Fig. 1A**, **Suppl. Fig. 2A**). Co-immunofluorescence analysis for desmin, one of the most reliable markers for HSCs in mouse liver (*33*) and GFP confirmed HSC-specific GFP expression (**Fig. 1B**). Primary HSCs isolated from *Pdgfrb*-GFP mice also co-expressed desmin and GFP (**Suppl. Fig. 2B**). We did not observe co-expression of GFP with markers of Kupffer cells (F4/80), liver sinusoidal endothelial cells (CD31/PECAM1) or hepatocytes (HNF4α) (**Suppl. Fig. 2C**) indicating that GFP is expressed only in HSCs.

**Fig. 1.**
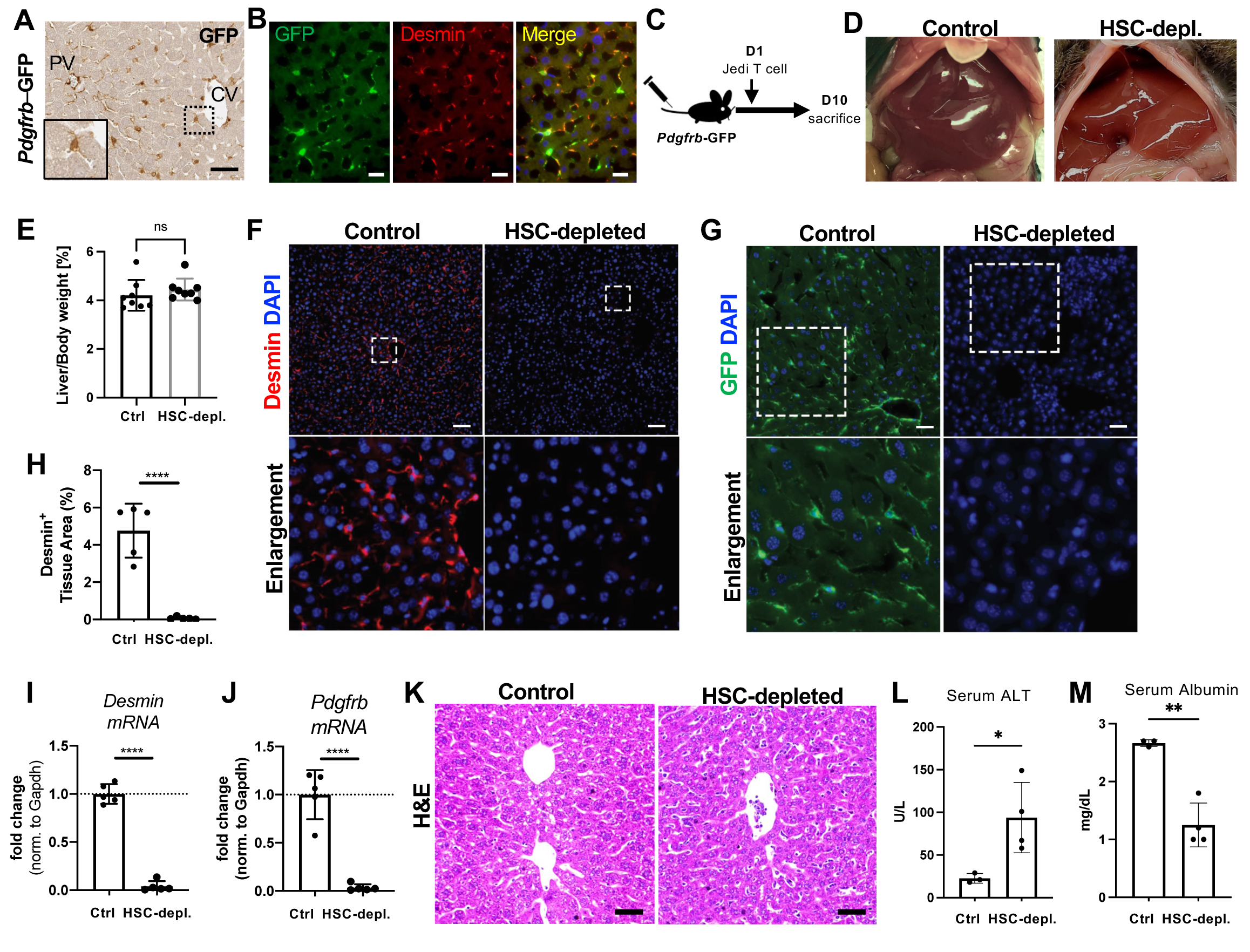
HSC depletion using Jedi T cells. **(A)** Immunostaining for GFP of *Pdgfrb*-GFP adult mouse liver. Scale bar represents 50 µm. **(B)** Immunofluorescence microscopy of liver section from *Pdgfrb*-GFP transgenic mice for GFP and desmin. Scale bars represent 20 µm. **(C)** Experiment outline. **(D)** Macroscopic liver images of control and HSC-depleted mice. **(E)** Liver to body weight ratios of control and HSC-depleted mice. Mean+SD. **(F)** Immunofluorescence microscopy of liver sections from control and HSC-depleted mice for desmin. Squares indicate areas of enlargement. Scale bars represent 100 µm. **(G)** Immunofluorescence microscopy of liver sections from control and HSC-depleted mice for GFP. Squares indicate areas of enlargement. Scale bars represent 50 µm. **(H)** Quantification of desmin positive tissue area in control and HSC-depleted mice. Mean+SD. **(I)** qRT-PCR for *Desmin* expression in whole liver RNA. Mean+SD**. (J)** qRT-PCR for *Pdgfrb* expression in whole liver RNA. Mean+SD. **(K)** H&E staining. Scale bars represent 50 µm. **(L)** Serum ALT levels. Mean+SD. **(M)** Serum albumin levels. Mean+SD. *p< 0.05, **p<0.01, ****p<0.0005. Two-tailed Student’s t-test.

Adoptive transfer of Jedi T cells isolated from Jedi mice, but not control T cells from H-2K^d^ positive control mice, almost completely ablated HSCs within 10 days in *Pdgfrb*-GFP mice (**Fig. 1C-J**). Livers appeared macroscopically normal, and HSC-depleted mice displayed a similar liver to body weight ratio as control mice (**Fig. 1D**, **E**). No desmin-, reelin-, GFAP- or GFP-positive cells could be detected by immunofluorescence, indicating efficient ablation of HSCs (**Fig. 1F**, **G**, **Suppl. Fig. 2D, E**). Quantification of the desmin-positive area by tissue morphometry revealed significantly fewer HSCs in HSC-ablated livers (**Fig. 1H**), a result that was confirmed by measuring *Desmin* and *Pdgfrb* gene expression in whole liver RNA by qRT-PCR (**Fig. 1I**, **J**). By H&E, hepatocytes appeared smaller, and sinusoids were slightly distended (**Fig. 1K**). Serum ALT levels were increased as an indication of discrete hepatocyte injury, and serum albumin levels decreased indicating reduced liver function in HSC-depleted mice (**Fig. 1L**, **M**). Of note, while Jedi T cells were readily detectable in the liver by granzyme B staining, a CD8 T cell marker, at 3 days after adoptive transfer, no Jedi T cells were detected at 10 days or up to 6 weeks after adoptive transfer (**Suppl. Fig. 3A**). Similarly, there was no Jedi *TCRa* or *TCRb* mRNA expression detected by whole liver RNA-seq at 10 days. These observations agree with earlier reports in which Jedi T cells become undetectable 2-3 weeks after adoptive transfer (*25, 26*).

In summary, these results demonstrate efficient ablation of HSCs with no detectable Jedi T cells following days after adoptive transfer in liver indicating that subsequent liver changes are due to ablation of HSCs rather than indirect effects of persistent Jedi T cells.

### Sustained HSC depletion causes loss of liver mass and diminished number of midlobular hepatocytes

The half-life of quiescent HSCs in mice is unknown. Therefore, we sacrificed mice 6 weeks after HSC ablation (**Fig. 2A**) and we found no evidence of desmin- or GFP-positive HSCs **(Fig. 2B**, **Suppl. Fig. 4A),** indicating that HSCs are completely depleted for up to 6 weeks. Unexpectedly, macroscopic inspection showed smaller livers in HSC-depleted mice (**Fig. 2C**), which was consistent with a significant decrease in liver to body weight ratio of up to ∼45% compared to controls (vehicle only injected mice) (**Fig. 2D**). While body weights were similar among both groups, liver weights were significantly lower in HSC-depleted mice (**Suppl. Fig. 4B, C**). H&E staining showed distended sinusoids and an increase in acellular areas reminiscent of large hepatic vessels (**Fig. 2E-F**, **Suppl. Fig. 4E**). However, based on immunostaining with the endothelial cell marker CD31/PECAM1, these structures were not lined by endothelial cells (**Fig. 2G**) and thus are acellular cavities. These cavities (annotated with blue circles) were consistently localized between central veins (yellow circles) and portal veins (green circles) within the midlobular zone 2 of the hepatic lobule. (**Fig. 2H**, **Suppl. Fig. 4E**).

**Fig. 2.**
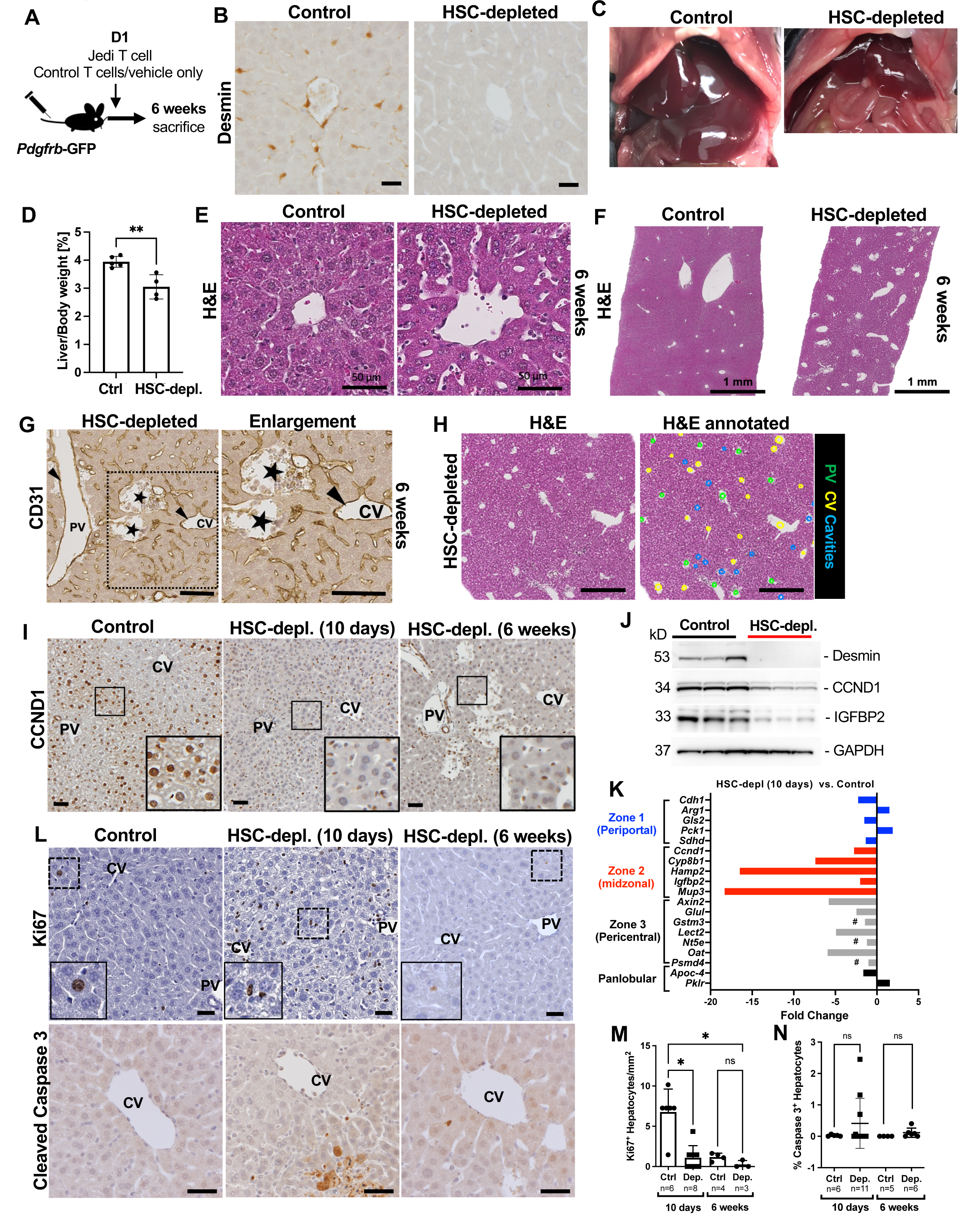
HSC depletion decreases liver mass and diminishes number of midlobular hepatocytes. **A)** Experimental outline. **(B)** Immunostaining for desmin of control and HSC-depleted mice 6 weeks after adoptive transfer. Scale bar represents 20 µm. **(C)** Macroscopic image of control and HSC-depleted mouse livers *in situ*. **(D)** Liver to body weight ratios of control and HSC-depleted mice at 6 weeks. **p<0.004 by unpaired two-tailed Student’s t-test. **(E)** H&E staining of control and HSC-depleted livers 6 weeks after adoptive transfer. Scale bars represent 50 µm. **(F)** Low magnification H&E staining of control and HSC-depleted livers 6 weeks after adoptive transfer. Scale bar represents 1 mm. **(G)** Immunostaining for CD31 of HSC-depleted liver sections. Arrows indicate endothelial lining of vessels. PV, portal vein, CV central vein, stars indicate acellular cavities with absent endothelial lining. Square indicates area of enlargement. Scale bars represent 100 µm. **(H)** Annotation of H&E staining of HSC-depleted liver at 6 weeks. Green dots indicate portal vein (PV), yellow dots central vein (CV), blue dots acellular cavities. Scale bar represents 500 µm. **(I)** Immunostaining for CCND1 of control and HSC-depleted livers at 10 days and 6 weeks after adoptive transfer. Squares indicate areas of enlargement in inserts. Scale bar indicates 50 µm. **(J)** Western blot of whole liver lysates from control and HSC-depleted livers 10 days after adoptive transfer for desmin, CCND1, IGFBP2, and GAPDH. **(K)** Whole liver RNA-seq analysis of control and HSC-depleted livers 10 days after adoptive transfer (n=6 controls vs. n=4 HSC-depleted mice). #, non-differentially expressed genes (FDR>0.34). **(L)** Immunostaining for Ki67 and cleaved caspase 3 of control and HSC-depleted livers at 10 days and 6 weeks after adoptive transfer. Scale bars represent 30 µm. Dashed rectangles indicate areas of enlargement in inserts. PV, portal vein. CV, central vein. **(M)** Quantification of Ki67-positive hepatocytes per mm^2^ in control and HSC-depleted livers at 10 days and 6 weeks after adoptive transfer. Mean+SD. Kruskal-Wallis’s test with post hoc Dunn’s test. *p<0.05 **(N)** Quantification of % cleaved caspase 3-positive hepatocytes per mm^2^ in control and HSC-depleted livers at 10 days and 6 weeks after adoptive transfer. Mean+SD. ns, not significant.

Two recent studies established the role of midlobular zone 2 hepatocytes in maintaining liver homeostasis and liver mass in healthy livers (*11, 12*). Central to this process is CCND1, that is constitutively expressed in most midlobular hepatocytes and critical for their proliferation (*11*). We thus investigated liver zonation and expression of CCND1 in HSC-depleted livers and control livers. As expected, control livers showed strong nuclear staining in midlobular zone 2 hepatocytes. By contrast, there was a markedly decreased nuclear hepatocyte CCND1 expression 10 days after HSC ablation which remained low for up to 6 weeks after HSC ablation (**Fig. 2I**). The rapid decrease in CCND1 was confirmed by western blotting of whole liver lysates for CCND1, as well as for another zone 2 hepatocyte marker, insulin growth factor-binding protein 2 (IGFBP2) (*11*) (**Fig. 2J**). Marked reduction in genes characteristic for zone 2 hepatocytes was also apparent by liver transcriptome analysis assessed at 10 days after HSC ablation. Based on whole liver RNA-seq analysis, there was a greater (up to -18.3-fold) and more consistent decrease in marker genes (*2, 11*) for zone 2 hepatocytes (*Hamp2*, *Cyp8b1*, *Mup3*, *Ccnd1* and *Igfbp2*) than in genes characteristic for zone 3 (*Axin2, Lect2, Oat, Nt5e, Glul*), or zone 1 (*Gls2, Arg1, Cdh1*), or panlobular genes (*Apoc4, Pklr*) (**Fig. 2K**, **Suppl. Fig. 4D**). While zone 3 genes *Axin2* and *Glul* were decreased (-5.8-fold and -2.5-fold, respectively, FDR<0.01), other zone 3 hepatocyte genes, *Gstm3*, *Nt5e* and *Psmd4,* were not differentially expressed (**Suppl. Fig. 4D**).

Gene expression analysis for metabolic pathways of HSC-depleted livers compared to control livers in KEGG, BioCarta and Reactome signatures (FDR<0.1) showed decreases in Wnt/β-catenin signaling, hedgehog signaling, and TGF-β1 signaling, and decreases in insulin receptor signaling, amino acid metabolism and cytochrome P450 metabolism some of which are to be expected, given the metabolic function of midlobular hepatocytes (*34*) and HSC-derived growth factor signaling pathways (*35*) (**Suppl. Table 1**). Stable expression of both glutamine synthetase, a pericentral marker and Wnt/β-catenin target (*36, 37*), and E-cadherin, a periportal marker, indicated that zone 1 and zone 3 hepatocytes were preserved both at 10 days and 6 weeks after HSC depletion (**Suppl. Fig. 4F, G**) despite decrease in Wnt/β-catenin signaling by gene expression analysis.

The liver’s size is tightly controlled and loss in liver mass – e.g., after partial liver resection – elicits a robust regenerative response. Thus, the loss in liver mass and decrease in midlobular CCND1 expression led us to investigate whether the regenerative response was impaired. Indeed, staining for Ki67 showed a lack of proliferating hepatocytes at either 10 days or 6 weeks after HSC depletion **(Fig. 2L**, **M).** Immunostaining for cleaved caspase 3, an apoptosis marker, showed positive hepatocyte staining in some mice though the increase was not significant and observed only in some HSC-depleted mice (n=3 out of 11 animals) (**Fig. 2L**, **N**).

In summary, our results suggest that HSCs are critical for maintaining midlobular hepatocyte CCND1 expression and liver mass.

### HSC-derived neurotrophin-3 induces CCND1 and is a hepatocyte mitogen

To investigate proteomic changes due to HSC ablation, we assessed whole liver protein lysates from control and HSC-depleted livers using the Olink mouse exploratory assay, which employs proximity extension assay technology for protein biomarker analysis of 92 proteins involved in diverse cellular processes (*38*). The Olink analysis separated control and HSC-depleted livers 10 days after depletion into two clearly distinct groups (**Fig. 3A**, **B**). In total, we found robust differences in 46 proteins between depleted and control livers, with levels of 30 proteins higher and 16 proteins lower in HSC-depleted than in control livers (adjusted P value <0.05). HGF was among the proteins expressed at lower levels, which was expected, given that it is produced and released by both HSCs and liver sinusoidal endothelial cells. Among the reduced proteins with a less well-established role in HSCs was neurotrophin-3 (NTF3), a member of the nerve growth factor family (**Fig. 3C**). HSCs are known to express both neuronal and glial genes such as glial fibrillary acidic protein (GFAP) and p75NTR/NGFR. Although it has been known for over 20 years that HSCs express NTF3 (*39*), its function in the liver remains largely unknown. We confirmed that NTF3 is expressed in HSCs by analyses of three published mouse and human liver scRNA-seq datasets (*3, 28, 30*), which showed highly specific *NTF3* expression in mesenchymal cells and HSCs (**Suppl. Fig. 4A-C**). Furthermore, our immunofluorescence analysis for NTF3 and desmin in mouse liver (**Fig. 3F**) and western blotting of mouse and human HSC cell lines JS1, LX2 and TWNT4 also showed robust NTF3 expression (**Fig. 3D**).

**Fig. 3.**
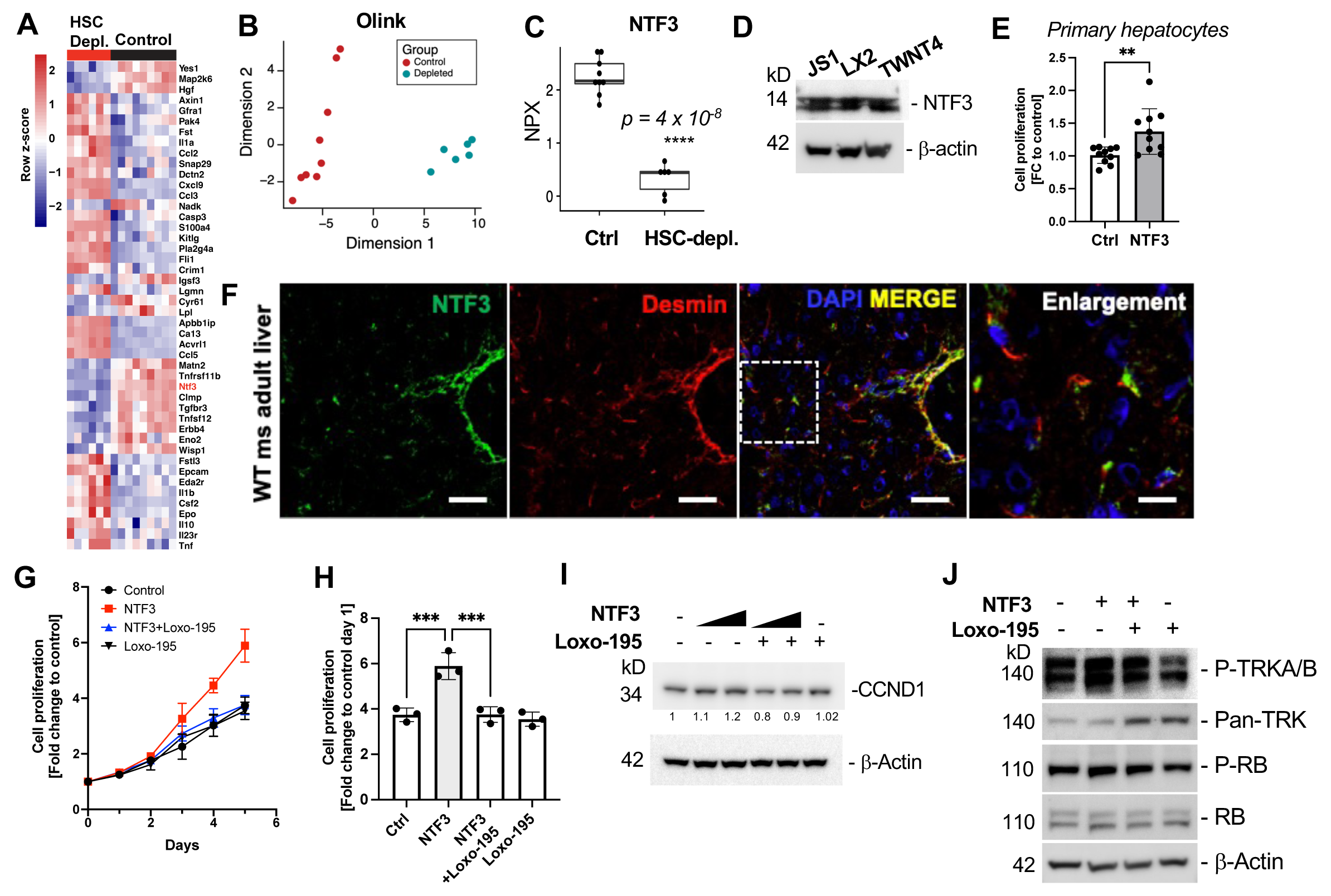
NTF3 is an HSC-derived hepatocyte mitogen that increases CCND1. **(A)** Protein biomarker analysis of control and HSC-depleted livers 10 days after adoptive transfer. **(B)** Multidimensional scaling of Olink biomarker analysis of whole liver lysates from control and HSC-depleted livers 10 days after adoptive transfer. **(C)** Comparative protein level analysis of NTF3 in control and HSC-depleted livers 10 days after adoptive transfer. **(D)** Western blot of mouse and human HSC cell lines for NTF3 and β-actin. (**E**) Cell proliferation of primary wildtype mouse hepatocytes after incubation with NTF3 (20 ng/ml) or vehicle only in serum-free William’s E media. Data from 2 independent experiments. Mean+SD. **p=0.0062 two-tailed Student’s t-test. **(F)** Immunofluorescence microscopy for NTF3 and desmin of wildtype adult mouse liver. Dashed rectangle indicates area of enlargement. **(G)** Cell proliferation analysis of HepG2 cells incubated with NTF3, NTF3/Loxo-195, Loxo-195 or vehicle only. Data from 3 independent experiments. Mean+SD. **(H)** HepG2 cell proliferation at 5 days incubated with NTF3, NTF3/Loxo-195, Loxo-195 or vehicle only. Data from 3 independent experiments. Mean+SD. ***p=0.009 by one-way ANOVA, post hoc Tukey’s test. **(I)** Western blot analysis for CCND1 and β-actin of cell lysates from HepG2 cells incubated with NTF3, NTF3/Loxo-195, Loxo-195 or vehicle only. **(J)** Western blot analysis of HepG2 cell lysates for Phospho-TRK (pTRKA^Y674/675^, pTRKB^Y706/707^), pan-TRK, Phospho-RB^S780^, RB and β-Actin.

The mature NTF3 protein is highly conserved in vertebrates. The human and mouse orthologs share 96% amino acid identity. We therefore incubated freshly isolated primary mouse hepatocytes in serum-free (**Fig. 3E)** and in FBS-containing media **(Suppl. Fig. 5H)** supplemented with recombinant NTF3 or vehicle control. We found that cell numbers were significantly higher in NTF3-treated primary hepatocytes than in control-treated hepatocytes **(Fig. 3E**, **Suppl. Fig. 5H)**. Similarly, NTF3 treatment of the mouse hepatocyte AML12 cell line (**Suppl. Fig. 5E, F**) and the human HepG2 cell line (**Fig. 3G**, **H**) greatly stimulated cell proliferation. NTF3 binds and activates tropomyosin receptor kinases (TRK) TRKA, TRKB, and TRKC, which are encoded by *NTRK1-3* genes (*40*). Binding of NTF3 to TRKs induces phosphorylation and activation of downstream signaling pathways (*40*). NTRK fusion-positive cancers have constitutive receptor activity and are oncogenic drivers, rendering them therapeutically responsive to TRK inhibitors (*41, 42*). We thus investigated whether NTF3-mediated cell proliferation is inhibited by the highly specific TRKA/TRKC inhibitor, Loxo-195 (selectritinib). Indeed, Loxo-195 (final concentration 5 nM) significantly inhibited cell proliferation in AML12 and HepG2 cells indicating that NTF3 induces cell proliferation by TRK signaling (**Fig. 3G-H**, **Suppl. Fig. 5D-E**). As HSC depletion decreases hepatocyte CCND1 expression **(Fig. 2I**, **J)**, we investigated if NTF3 stabilizes CCND1 in culture. Incubation of HepG2 with NTF3 increased CCND1 expression, which was inhibited by the addition of Loxo-195 (**Fig. 3I**). We assessed activation of downstream pathways of both TRK and CCND1 signaling, which demonstrated increased TRK phosphorylation (P-TRKA^Tyr674/675^, P-TRKB ^Tyr706/707^) and increased phosphorylation of retinoblastoma (*RB1*), respectively, which were both decreased by addition of Loxo-195, supporting the conclusion that NTF3 stabilizes CCND1 and activates CCND1 downstream pathways via TRK activation (**Fig. 3J**). These results were consistent when using NTF3 from three different vendors was used; in all cases, CCND1 levels increased in both AML12 and HepG2 cells (**Suppl. Fig. 5F, G**).

### Administration of recombinant neurotrophin-3 to HSC-depleted mice rescues CCND1 expression and increases liver mass

In our murine model, ablation of HSCs decreased nuclear CCND1 in midlobular hepatocytes (**Fig. 2I**, **J**). To determine whether NTF3 is required for midlobular CCND1 expression, we assessed if *in vivo* administration of NTF3 increased hepatocyte CCND1 expression in HSC-depleted mice. We induced HSC depletion and injected mice with NTF3 at two concentrations, 20 ng/g and 100 ng/g body weight, or vehicle control for 4 consecutive days. Mice were sacrificed after the last injection (**Fig. 4A**, **B**). Liver to body weight ratio of mice treated with NTF3 was dose-dependently increased with a significant increase in liver to body weight ratio in the higher dose group (**Fig. 4C**). As expected, in HSC-depleted mice treated with vehicle only, CCND1 was decreased in hepatocytes and there was some scattered CCND1 expression in non-parenchymal cells (**Fig. 4D**). NTF3 treatment of HSC-depleted mice led to the reconstitution of nuclear CCND1 expression in hepatocytes in the midlobular region (**Fig. 4D**). To quantify CCND1 expression in hepatocytes, we assessed livers by multiplex immunohistochemistry for CCND1 and HNF4α, a hepatocyte marker, which is ubiquitously expressed in the nuclei of hepatocytes across the liver lobule in both control and HSC-depleted mice (**Suppl. Fig. 6A**).

**Fig. 4.**
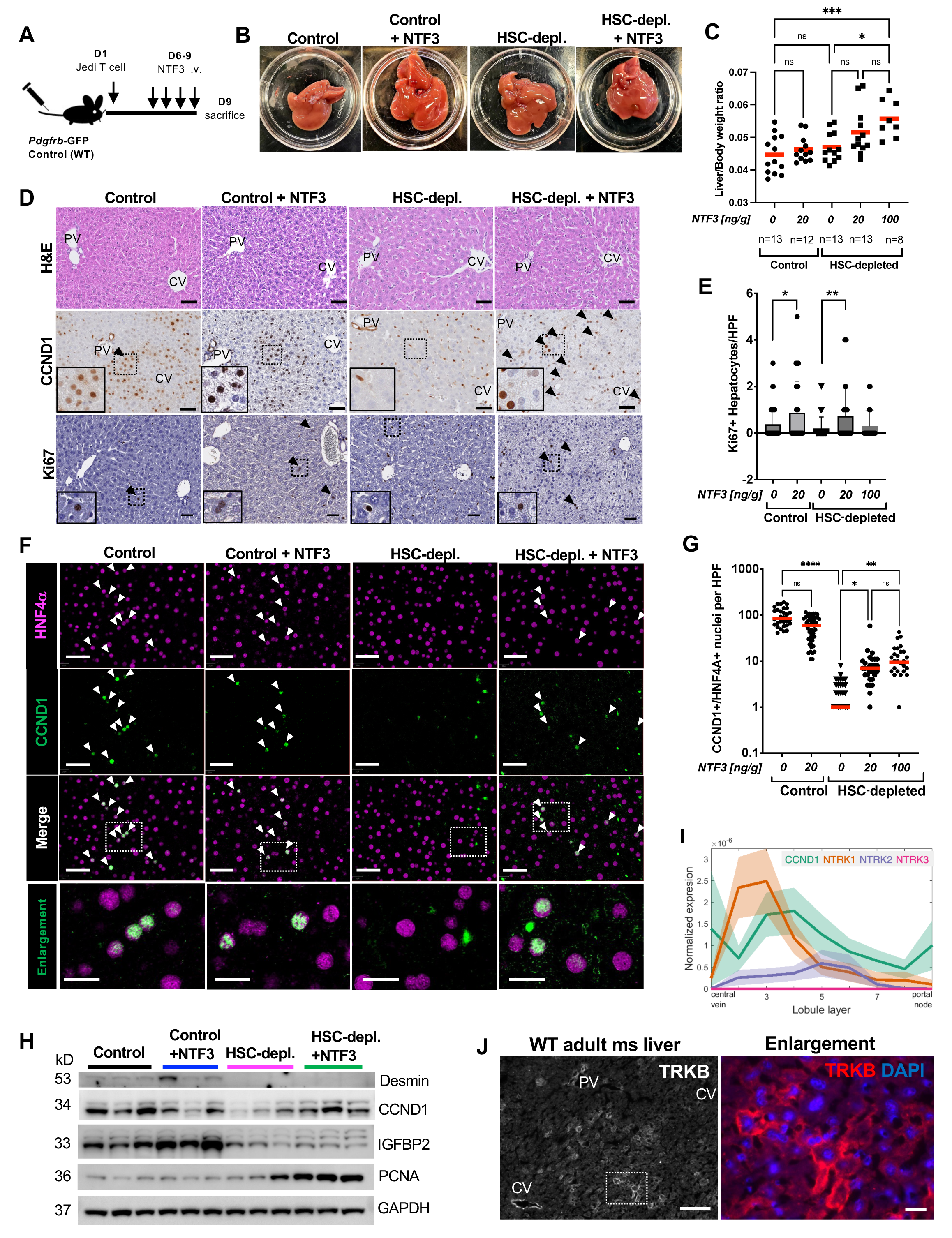
Recombinant NTF3 increases hepatocyte CCND1 expression in HSC-depleted mice. **(A)** Experimental outline. **(B)** Macroscopic liver images of controls and HSC-depleted mice treated with vehicle or NTF3. **(C)** Liver to body weight ratios of controls and HSC-depleted mice treated with vehicle or NTF3. *p=0.0184, ***p=0.0005 by one-way ANOVA, post hoc Tukey’s test. **(D)** H&E, CCND1 and Ki67 staining of liver sections from control and HSC-depleted mice treated with vehicle or NTF3. Arrows indicate positive staining. Dashed rectangles indicate areas of enlargement. Scale bars represent 40 µm. **(E)** Quantification of Ki67-positive hepatocytes in control and HSC-depleted mice treated with vehicle or NTF3. Mean+SD. Two-tailed Student’s t-test. *p=0.0235, **p=0.008 **(F)** Multiplex immunohistochemistry for CCND1 and HNF4α of liver sections from control and HSC-depleted mice treated with vehicle or NTF3α Dashed rectangles indicate areas of enlargement. Scale bars represent 20 µm. Arrows indicate positive-staining nuclei. **(G)** Quantification of CCND1/HNF4α double-positive nuclei per high power field (HPF). Bars indicate mean. Kruskal-Wallis, post hoc Dunn’s test. *p=0.028, **p=0.0048, ****p<0.0001. **(H)** Western blotting for desmin, CCND1, IGFBP2, PCNA, GAPDH of whole liver lysates from control and HSC-depleted mice treated with vehicle or recombinant NTF3. **(I)** Analysis of a published mouse liver gene expression dataset (*2*) which enables layer-specific hepatocyte gene expression analysis according to their location in the lobule layer for *CCND1, NTRK1, NTRK2, NTRK3*. **(J)** Immunofluorescence microscopy for TRKB (*NTRK2*) in normal adult mouse liver at 4 months of age. Dashed rectangle indicates area of enlargement. CV, central vein. PV, portal vein.

Quantification of double-positive CCND1 and HNF4α nuclei showed that treatment with NTF3 dose-dependently increased the number of double-positive nuclei (**Fig. 4F**, **G**). Similarly, we observed an increase in Ki67-positive hepatocytes consistent with increased regenerative activity in both HSC-depleted mice and in NTF3-treated control mice (**Fig. 4D**, **E**). Western blotting of whole liver lysates also confirmed that NTF3 treatment increased CCND1 expression and PCNA levels in HSC-depleted mice, further indicating increased regenerative activity (**Fig. 4H**).

Analysis of a published mouse liver dataset that allows for zone-specific gene expression analysis of hepatocytes (lobule layers 1 through 9) (*2*) showed gene expression of *NTRK1* and *NTRK2* in midlobular hepatocyte layers 2 through 7 (**Fig. 4I**) whereas *NTRK3* was absent.

Immunofluorescence analysis for TRKA (*NTRK1*) showed non-parenchymal TRKA staining around portal tracts and central veins **(Suppl. Fig. 6B).** TRKB (*NTRK2*) staining was observed in midlobular hepatocytes with decreased staining around portal tracts and central veins, suggesting that TRKB mediates NTF3 effects though further studies are required (**Fig. 4J**).

**Interpretations** HSCs provide mitogenic cues for zone 2 hepatocyte proliferation by releasing NTF3 to maintain liver homeostasis.

## Discussion

We have put forth a highly efficient HSC ablation model which enables us to uncover the role of quiescent HSCs in adult mouse livers during homeostasis. Previous HSC ablation models using diptheria-toxin receptor transgenic mice (*43*), thymidine kinase transgenic mice (*44*) or HSC ablation by gliotoxin administration (*45*) achieved only partial (50-65%) ablation of HSCs. While we have focused on the HSC-hepatocyte interaction, this ablation model might also be useful for investigating the interaction of HSCs with other liver cells, e.g., the effects of HSCs on liver sinusoidal endothelial cell biology or how HSCs modulate the immune microenvironment (*46*). While GFP is highly expressed in HSCs within the liver, extrahepatic expression of GFP in PDGFRB-positive cells could be used to study pericyte biology in other organ systems.

Signaling pathways that regulate liver zonation have been best characterized for zone 1 and zone 3 (*1, 47–50*), which was facilitated by the longstanding tradition of binary classification of hepatocytes and well-established markers for both zone 1 (e.g. E-cadherin) and zone 3 (e.g. glutamine synthetase). Much less is known about signals determining midlobular hepatocyte zonation (*16*). Unique genes expressed at higher levels in midlobular hepatocytes have only been more recently identified using advanced scRNA-seq/spatial transcriptomics techniques and fate-labeling methods (*2, 11, 12, 34, 51*). These include hepcidin antimicrobial peptide *(Hamp2)* and *Igfbp2* (*2, 5, 11*) suggesting that midlobular hepatocytes play important roles in iron metabolism and insulin growth factor signaling. Midlobular hepatocytes also express higher levels of cytochrome P450 xenobiotic metabolism genes (*2, 34, 51*). Genetic fate-labeling in *Hamp2-CreER* transgenic mice confirmed *Hamp2* as a primarily midlobular marker (*11*) and identified IGFBP2-mTOR-CCND1 signaling as a critical pathway to maintain midlobular hepatocyte proliferation. Deletion of either *Igfbp2* or inhibition of mTORC1/2 led to decreased CCND1 expression and hepatocyte proliferation *in vivo* (*11*). While our results demonstrate that HSCs are essential to maintain midlobular CCND1 expression and liver homeostasis, NTF3 only increased CCND1 but not IGFBP2 expression in hepatocytes of HSC-ablated mice suggesting that NTF3-TRKB signaling may regulate CCND1 separately from the IGFBP2-mTOR-CCND1 axis *in vivo*.

As mentioned, fate-labeling studies indicated that midlobular hepatocytes have a higher proliferative capacity than periportal or perivenous hepatocytes in homeostatic liver (*11, 12*). The niche signaling determining this process was unknown (*16*). Our results indicate that HSC-derived NTF3 acts as a full-fledged hepatocyte mitogen. In liver regeneration, signals inducing hepatocyte proliferation may be differentiated into ‘complete mitogens’ and ‘auxiliary hepatocyte mitogens’ depending on their ability (1) to induce proliferation in cultured primary hepatocytes and (2) to induce hepatocyte DNA synthesis and liver enlargement *in vivo* (*15*). In contrast to complete mitogens, auxiliary mitogens cannot induce hepatocyte proliferation in culture or if injected into rodents. While auxiliary mitogens include several different molecules (*15, 52*), complete mitogens only include hepatocyte growth factor (HGF), and ligands of epidermal growth factor receptor (EGFR). Other complex signaling pathways can also promote hepatocyte proliferation but disruption of these, e.g., Wnt/β-catenin and hedgehog signaling – delays liver regeneration but do not abolish regeneration (*53, 54*). Although the role of NTF3 in liver regeneration remains to be clarified, our results demonstrate that NTF3 increases proliferation of cultured primary hepatocytes, and induces hepatocyte proliferation, and enlargement of liver size if injected into mice (**Fig. 4C**). Of note, NTF3 induced hepatocyte proliferation in HSC-depleted mice despite decreased protein levels of HGF, underscoring its effectiveness (**Fig. 3A**).

At this point, two question arise: how NTF3 signals *in vivo* to hepatocytes and why it might have the greatest effect on midlobular hepatocytes. Analysis of a published mouse liver scRNA-seq dataset that allows for expression analysis of hepatocytes according to their localization in the liver lobule (*3*) indicated that *NTRK1* and *NTRK2,* but not *NTRK3,* were expressed in midlobular hepatocytes (**Fig. 4I).** Immunofluorescence staining for TRKB (*NTRK2*) showed TRKB-positive midlobular hepatocytes, possibly explaining why NTF3 has the largest effect on midlobular hepatocytes; however, further studies characterizing TRK expression in the liver are required.

Overall, *NTRK* gene expression levels seem very low in liver in queries of publicly accessible datasets such as TCGA, Gtex, and our own bulk RNA-seq dataset of control and HSC-depleted livers. However, low expression of NTRKs in healthy livers might not preclude high specific receptor activity as exemplified by the efficiency of HER2-targeting antibody drug conjugates in breast cancers with low HER2 expression (*55*). Furthermore, besides hepatocytes, other cells – e.g., HSCs themselves – express high levels of *NTRK2*, *NTRK3,* and *NGFR* (*39, 56*). Thus, autocrine neurotrophin signaling in HSCs could be biologically relevant. HSCs are highly pleiotropic, therefore, other signaling pathways besides NTF3-TRKB might also be affected and contribute to the phenotype observed in HSC-depleted livers. Furthermore, while we have focused on how NTF3 affects CCND1, NTF3 might also affect other proteins and signaling pathways in either HSC-depleted or control livers.

In summary, HSCs have important physiological functions, and their role in liver homeostasis must be further elucidated if we are to deepen our understanding of pathobiological processes. While HSCs are the principal fibrogenic cells in chronic liver injury driving fibrosis towards end-stage liver disease, therapeutic strategies targeting HSCs in liver cirrhosis analogous to T cell-targeted immunotherapy for cardiac fibrosis, should be developed carefully and investigated in detail (*57–59*). NTF3 is a complete hepatocyte mitogen whose signaling pathway is well characterized in the nervous system and in cancer and for which signaling pathway agonists and antagonists (e. g., TRK inhibitors) are available (*41*). Future studies may well reveal how these pathways could be exploited therapeutically to e. g., promote liver regeneration and/or inhibit proliferation in hepatic malignancies.

## Acknowledgments

We thank Andrea Branch for critical reading and constructive feedback on the manuscript. Joseph Roland for expert advice and support by the DSHR at Vanderbilt University Medical Center, the Genomics Core at Rockefeller University for excellent support, Seung-Hee Kim-Schultze for expert support at the HIMC at Icahn School of Medicine at Mount Sinai, and the Department of Animal Care at Vanderbilt University Medical Center. Core Services (Flow core, DHSR) were performed through Vanderbilt University Medical Center’s Digestive Disease Research Center supported by NIH grant P30DK058404.

## Author contributions

YAL, JA, BB, SLF conceived the initial project. YAL, VQT, TFL, KR, MCY, SL, MCY, TT, JA, JC performed experiments. Olink and RNA-seq data analysis was performed by KMA. VQT, TL, KMA and YAL wrote the original draft. YAL, SLF acquired funding.

## Data and materials availability

The Gene Expression Omnibus accession number for the liver transcriptome profiles of HSC-depleted livers vs. controls reported in this paper is GSE211370. Analysis interface of published scRNA-seq datasets (*2, 3, 28, 30, 31*) shown for this paper are available here: https://shiny.igmm.ed.ac.uk/livermesenchyme/ (Dobie et al. Cell Reports, 2019), https://shiny.igmm.ed.ac.uk/livercellatlas/ (Ramachandran et al. Nature, 2019), https://itzkovitzwebapps.weizmann.ac.il/webapps/home/session.html?app=HumanLiverBrowser (Halpern et al. Nature, 2017; Massalha et al. Mol Systems Biol, 2020). https://itzkovitzapapp.weizmann.ac.il/apap/?gene1=Col1a1 (Ben-Moshe et al. Cell Stem Cell, 2022).

Remaining data is available on request by the corresponding author.

## Figure Legends

**Suppl. Fig. 1.**
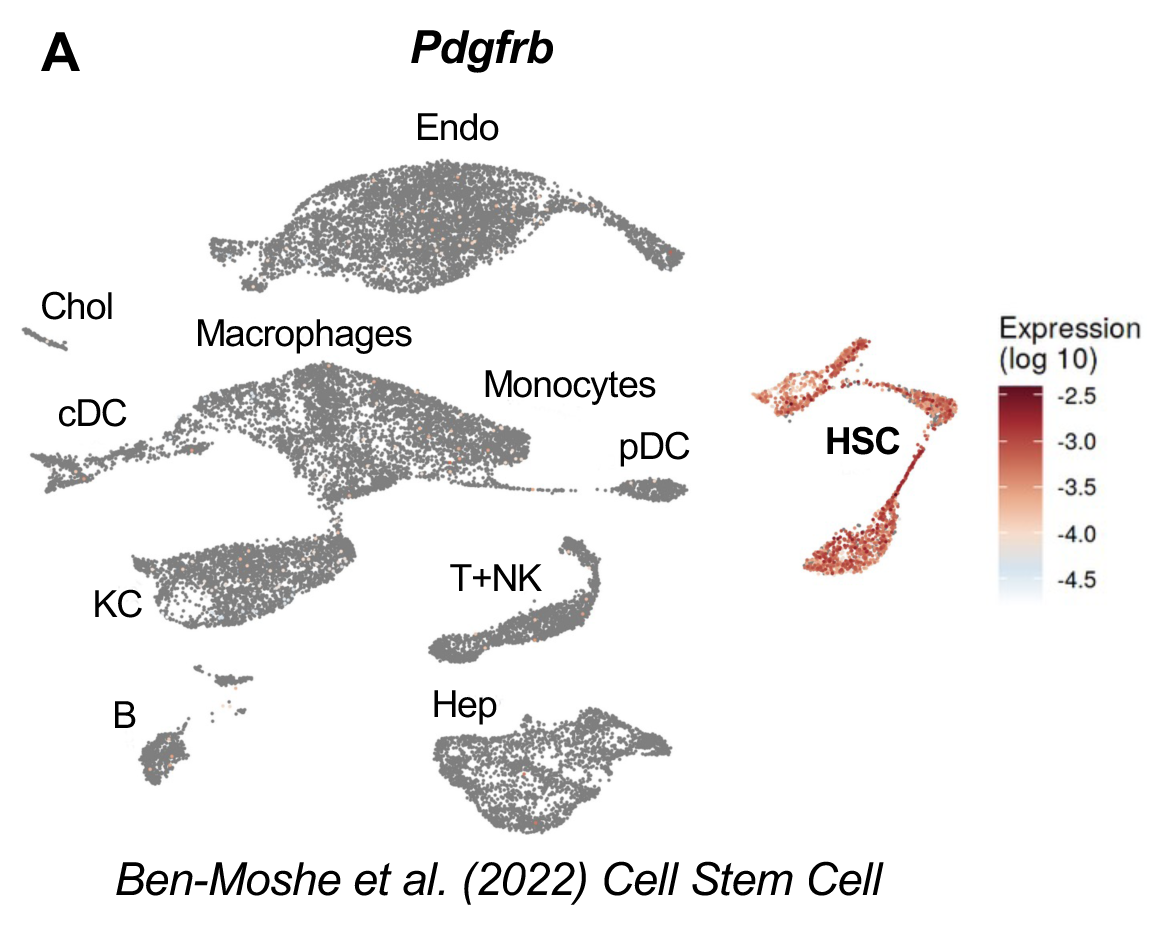
*Pdgfrb* is highly expressed by HSCs. **(A)** Analysis of a published mouse liver scRNA-seq dataset (*31*) from uninjured and acetaminophen-treated livers for *Pdgfrb*.

**Suppl. Fig. 2.**
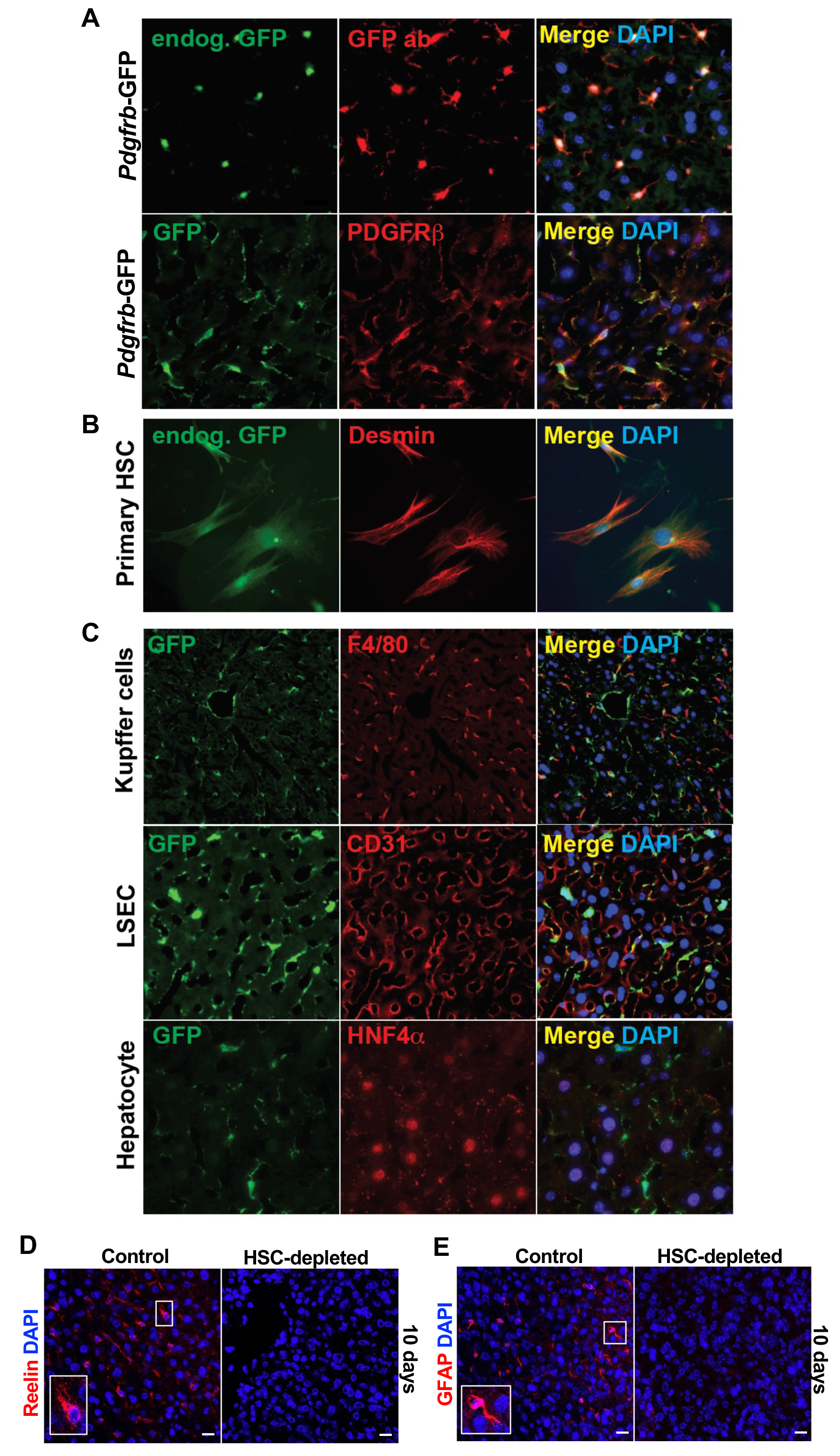
*Pdgfrb*-GFP transgenic mice selectively express GFP in HSCs within liver. **(A)** Immunofluorescence microscopy of liver sections from *Pdgfrb*-GFP mice for endogenous GFP fluorescence and anti-GFP antibody enhanced detection for GFP and PDGFRB. (B) Immunocytochemistry for GFP and desmin of primary HSCs isolated from *Pdgfrb*-GFP mice. (C) Immunofluorescence microscopy for GFP and F4/80, CD31 and HNF4α of liver sections from *Pdgfrb*-GFP mice. (D) Immunofluorescence microscopy of liver sections from control and HSC-depleted livers for reelin. Scale bars represent 200 µm. (E) Immunofluorescence microscopy of liver sections from control and HSC-depleted livers for GFAP. Squares indicate areas of enlargements (insert). Scale bars represent 200 µm.

**Suppl. Fig. 3.**
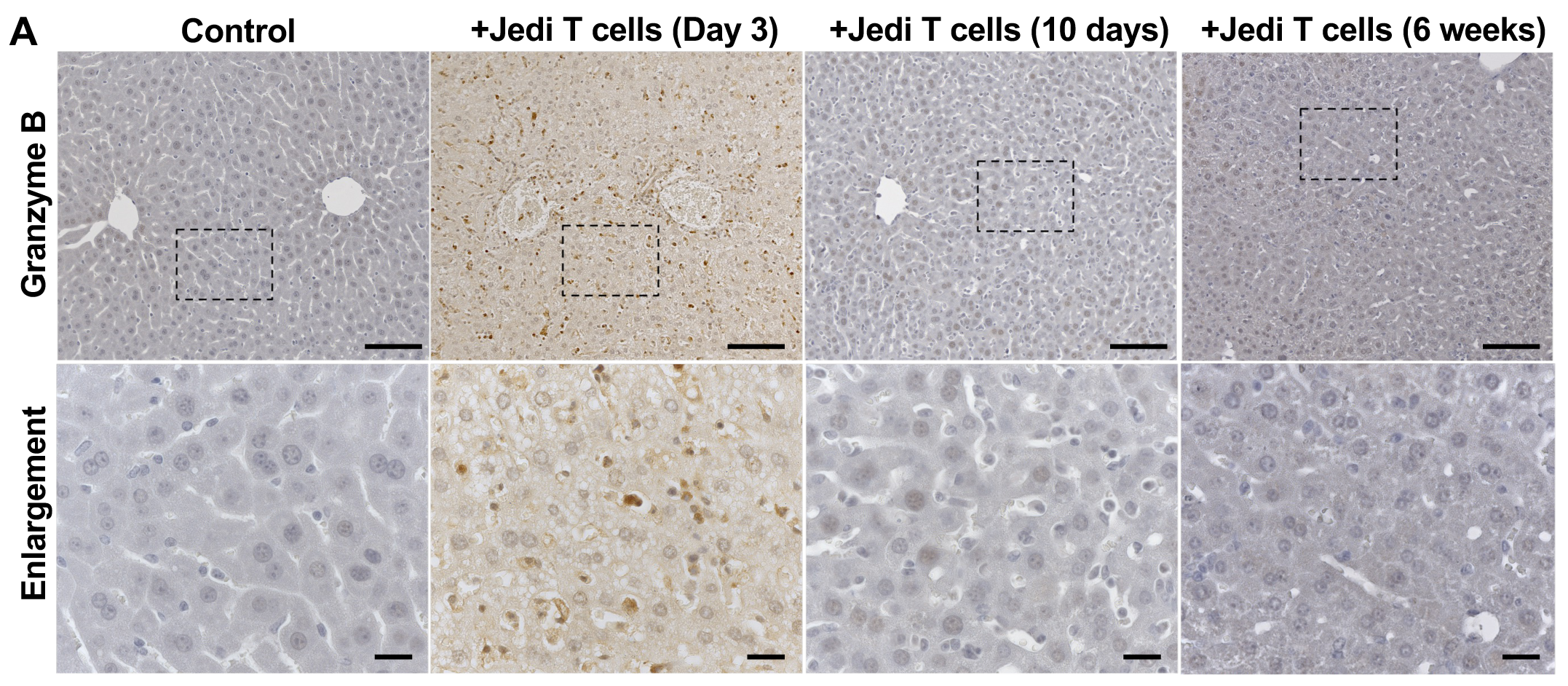
Time course of hepatic Jedi T cell expansion and clearance at 3 days, 10 days and 6 weeks after adoptive transfer in *Pdgfrb*-GFP mice. **(A)** Immunostaining for granzyme B of liver sections from control and *Pdgfrb*-GFP mice 3 days, 10 days, and 6 weeks after adoptive transfer. Dashed rectangles indicate areas of enlargement. Scale bars represent 100 µm (upper panel) and 20 µm (lower panel).

**Suppl. Fig. 4.**
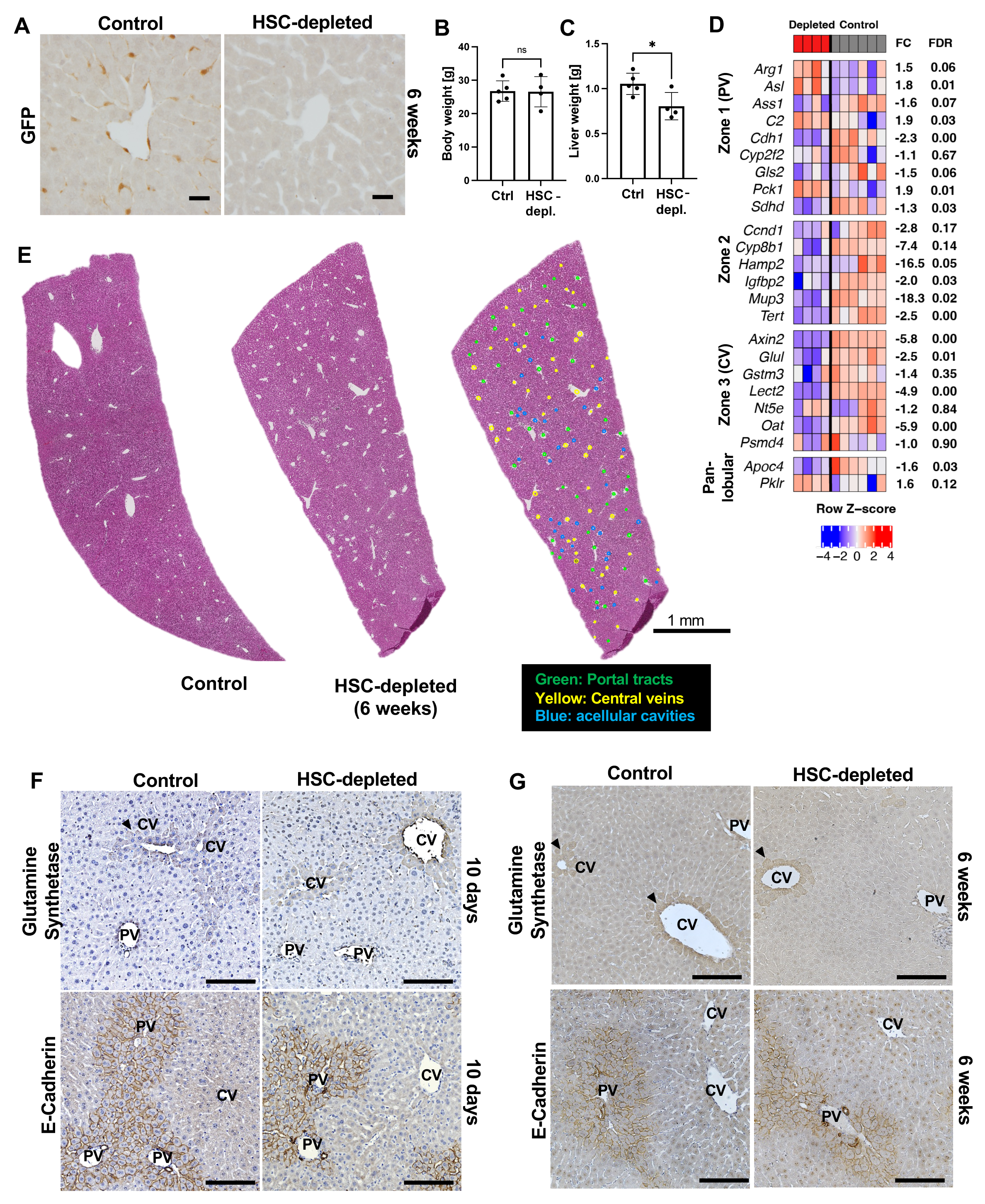
Diminished number of midlobular hepatocytes after sustained HSC depletion. **(A)** Immunostaining for GFP of control and HSC-depleted livers 6 weeks after adoptive transfer. Scale bar represents 20 µm. **(B)** Body weights of control and HSC-depleted mice 6 weeks after adoptive transfer. Mean+SD. **(C)** Liver weights of control and HSC-depleted mice 6 weeks after adoptive transfer. Mean+SD. *p=0.02 unpaired two-tailed t-test. **(D)** Gene expression analysis of whole liver RNA from HSC-depleted mice vs. controls 10 days after adoptive transfer. FC, fold change; FDR, false discovery rate. (**E)** Low magnification and annotation of H&E staining of control and HSC-depleted livers 6 weeks after adoptive transfer. Scale bar represents 1 mm. **(F)** Immunostaining for glutamine synthetase and E-cadherin of control and HSC-depleted livers 10 days after adoptive transfer. Arrows indicate areas of positive staining around CVs. Scale bars represent 50 µm. **(G)** Immunostaining for glutamine synthetase and E-cadherin of control and HSC-depleted livers 6 weeks after adoptive transfer. Arrows indicate areas of positive staining around CVs. Scale bars represent 50 µm. CV, central vein. PV, portal vein.

**Suppl. Fig. 5.**
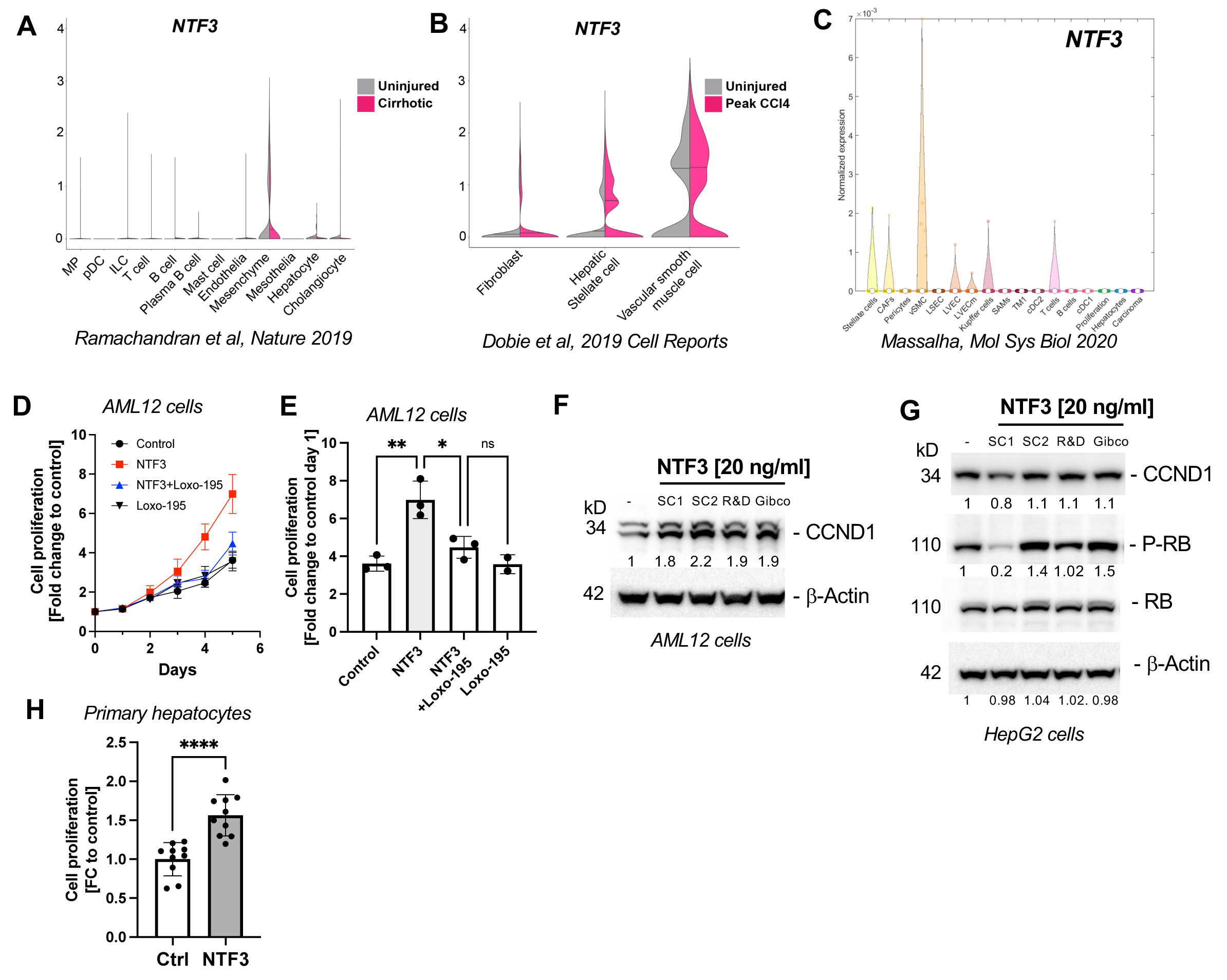
NTF3 is an HSC-derived hepatocyte mitogen. **(A)** Analysis of a published scRNA-seq dataset (*30*) of uninjured and cirrhotic human livers for *NTF3*. **(B)** Analysis of a published scRNA-seq dataset (*28*) of uninjured and CCl_4_-treated mouse livers for *NTF3*. **(C)** Analysis of a published scRNA-seq dataset (*3*) of normal human liver and hepatocellular carcinoma for *NTF3*. **(D**) Cell proliferation analysis of murine AML12 cells incubated with NTF3, NTF3/Loxo-195, Loxo-195 or vehicle only. Mean+SD. Data from 3 independent experiments. Mean+SD. **(E)** Cell proliferation of AML12 cells incubated with NTF3, NTF3/Loxo-195, Loxo-195 or vehicle only at day 5. Mean+SD. **p=0.002, *p=0.01, one-way ANOVA, post hoc Tukey’s test. **(F)** Western blotting for CCND1 and β-Actin of AML12 cells incubated with recombinant NTF3 from three different vendors. **(G)** Western blotting for CCND1, Phospho-RB^S780^, RB and β-Actin of cell lysates from HepG2 cells incubated with recombinant NTF3 from three different vendors. (**H**) Cell proliferation of primary wildtype mouse hepatocytes after incubation with NTF3 (20 ng/ml) or vehicle only in DMEM/5% FBS. Data from 2 independent experiments. Mean+SD. ****p<0.0001 by unpaired, two-tailed Student’s t-test.

**Suppl. Fig. 6.**
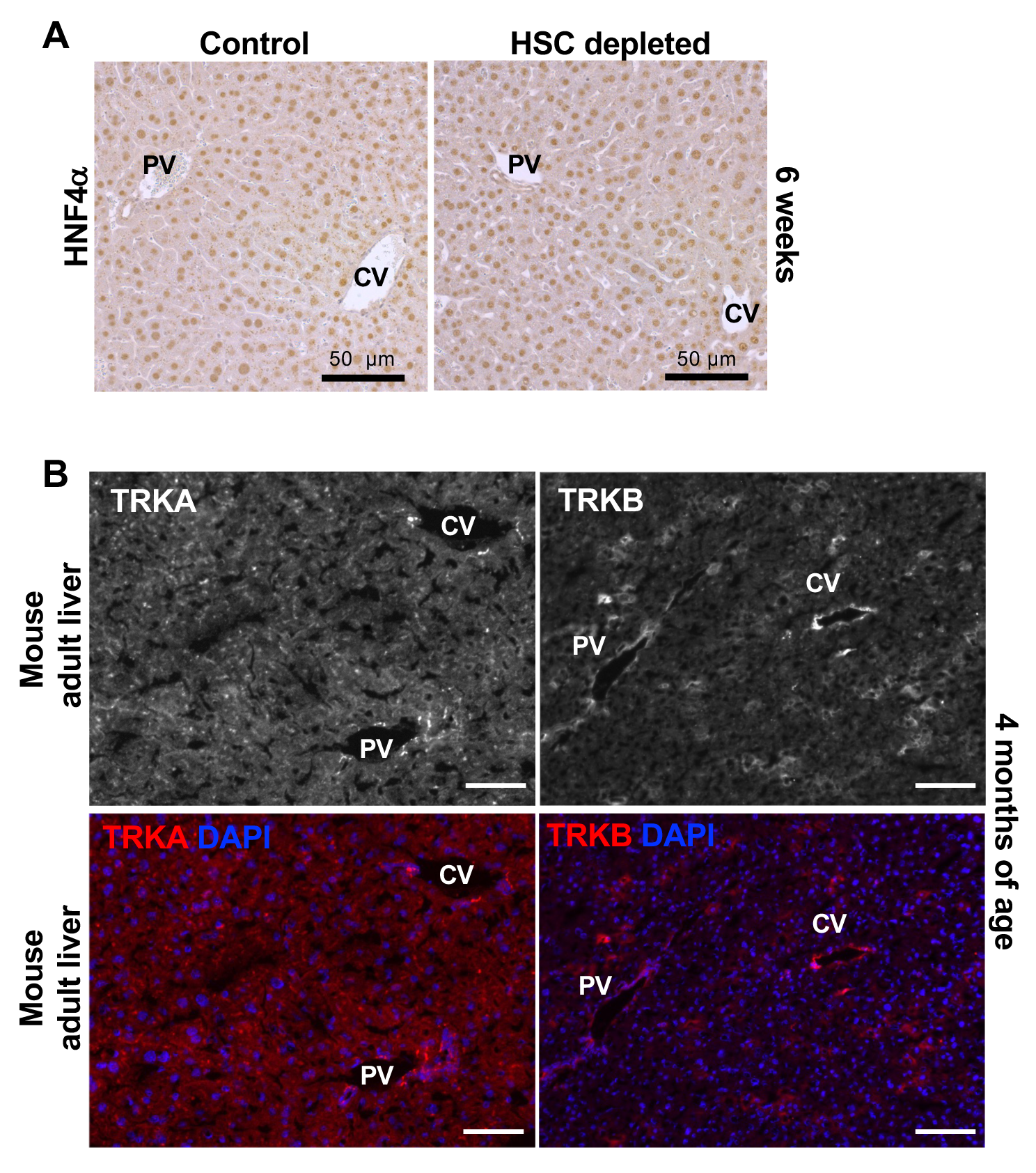
Pan-lobular expression of HNF4α in hepatocytes of control and HSC-depleted mice. **(A)** Immunostaining for HNF4α of control and HSC-depleted livers 6 weeks after adoptive transfer. Scale bar represents 50 µm. **(B)** Immunofluorescence microscopy for TRKA (*NTRK1*) and TRKB (*NTRK2*) of adult mouse liver. PV, portal vein. CV, central vein.

## Supplementary Materials for

### Materials and Methods

#### Mice

Jedi mice were obtained from Jackson Laboratories (Jax 028062) and are homozygous for CD45.1 and H-2K^d^ haplotype. *Pdgfrb*-BAC-eGFP cryopreserved sperm was obtained from MMRRC (031796-UCD), and mice were rederived in C57Bl/6, sub strain B10.D2 females (Jax 000463) that express CD45.2 and H-2K^d^ haplotype. Rederived *Pdgfrb*-GFP mice were backcrossed into B10.D2 and were maintained heterozygous for GFP, homozygous for CD45.2 and homozygous for H-2K^d^ haplotype. All experiments were performed when mice were 6 weeks to 6 months old and both male and females were used. All animal procedures were performed according to protocols approved by the Institutional Animal Care and Use Committees at Icahn School of Medicine at Mount Sinai and the Institutional Animal Care and Use Committees at Vanderbilt University Medical Center.

#### Genotyping

Jedi mice were genotyped by PCR genotyping using the following primers: TCRα-For: GAGGAGCCAGCAGAAGGT; TCRα-Rev: TCCCACCCTACACTCACTACA; TCRα-For: TCAAGTCGCTTCCAACCTCAA; TCRα-Rev: TGTCACAGTGAGCCGGGTG. *Pdgfrb*-GFP mice were genotyped by PCR using the following primers: GFP-For: GTGGAAGCAGAGAGGAGAGCATTTG, GFP-Rev: GGTCGGGGT-AGCGGCTGAA. H-2K^d^ haplotype, CD45.1, CD45.2 status were determined by flow analysis of PBMCs using the following antibodies purchased from ebiosciences: CD45.1-eFluor-450 (48-0453-80), CD45.2-PE (12-0454-82), H-2K^d^-APC (17-5957-80).

#### T cell isolation and adoptive transfer

CD8 T cells were isolated from spleen and lymph nodes (brachial, inguinal, axillar, and mesenteric) of Jedi and controls. Spleens and lymph nodes were mechanically homogenized and filtered through a 70 µm cell strainer (Falcon). After RBC lysis (Thermoscientific, A10492), CD8 T cells were enriched using the mouse CD8 T cell isolation kit (STEMCELL technologies 19853A) according to the manufacturer’s instructions. Mice with less than 25 g body weight were injected with up to 12 million Jedi T cells and mice greater than 25 g body weight were injected by tail vein injection with 18 million Jedi T cells. Control mice were injected with either T cells from H-2K^d^ mice or with vehicle only. Mice were not injected with lentiviral GFP particles as previously described (*1, 2*) to avoid lentiviral GFP-expression of other cells such as hepatocytes.

#### Treatment with recombinant NTF3

Four months old mice were injected via tail vein for 4 consecutive days with recombinant NTF3 (STEMCELL technology, cat. 78074, lots 1000070683, 1000077434) diluted in sterile 0.9% saline (0.9% NaCl) or vehicle only. Mice were weighed daily and recombinant NTF3 given at 20 ng/g body weight or 100 ng/g body weight, respectively. Lyophilized NTF3 was solubilized in 0.1% BSA/PBS and stored at -80°C until use. As activity and protein concentration declined significantly at approximately 3 months, only aliquots stored for 3 months or less were used for experiments.

#### Mouse blood chemistry analysis

Mouse serum or EDTA-plasma were stored at -80°C after collection and analyzed on a VetScan vs2 (Abaxis) using mammalian liver profile analysis discs (Abaxis, 500-0040) according to the manufacturer’s instructions.

#### RNA extraction and qRT-PCR

Bulk liver total RNA was isolated using the RNeasy Kit (Qiagen) following the manufacturer’s instructions. Five µg of total RNA was transcribed to cDNA using the RNA-to-cDNA ecoDry Premix (Takara). IQ SYBR Green Supermix (Biorad) was used to quantitative on a LightCycler480 System (Roche) or CFX96 Real-time System (Biorad). Primer sequences used were *Desmin*: For-GCCACCATGAGCCAGGCCTACT, Rev-TGCTCGAGGGAAC-ACGGGAGA, *Pdgfrb*: For-ACTACATCTCCAAAGGCAGCACCT, Rev-TGTAGAACTGGTCGTTCATGGGCA, *GAPDH:* For-CAATGACCCCTTCATTGACC, Rev-GATCTCGCTCCTGGAAGATG.

#### RNA-sequencing

Bulk total RNA was isolated using RNeasy Kit (Qiagen) following the manufacturer’s instructions, and 1 µg were used for polyA selection and library preparation using the Illumina TruSeq protocol. Samples were sequenced 50 bp paired-end on a NovaSeq 6000 by the Genomics Core Facility at The Rockefeller University. Raw FASTQ files were aligned to the mouse mm10 genome using the STAR aligner (v. 2.7.1a). The STAR command used was

~~~
STAR --runThreadN 10 \\
      --genomeDir mm10 \\
      --readFilesIn INPUT_R1_001.fastq.gz INPUT_L001_R2_001.fastq.gz \\
      --clip3pAdapterSeq GATCGGAAGAGC \\
      --outFilterMismatchNmax 2 \\
      --outFileNamePrefix OUTFILE \\
      --readFilesCommand gunzip -c \\
      --outSAMunmapped Within \\
      --outStd BAM_Unsorted \\
      --outSAMtype BAM Unsorted \\
      --outStd BAM_Unsorted
~~~

The resulting BAM file was sorted using samtools (v. 1.15.1). Reads were summarized using featureCounts (v1.6.4) against the Gencode mouse vM6 release disregarding multi-mapping and overlapping reads. The command used was:

~~~
featureCounts -a INPUT.gtf -T 4 -s 2 -o OUTFILE.txt LIST_OF_BAMFILES
~~~

Differential and gene set analysis was done in R session (development version, 2022-01-07 r81454) using the quasi-likelihood framework in edgeR (v. 3.37.0) and the mroast function on the mouse version of the MSigDB gene sets based on release v5.2 provided by the Bioinformatics group at the Walter+Eliza Hall Institute for Medical Research (https://bioinf.wehi.edu.au/software/MSigDB/). For other packages used to generate figures see section *Olink*. The Gene Expression Omnibus accession number for the transcriptome profiles reported in this paper is GSE211370.

#### Olink

Fresh frozen liver tissue samples were homogenized using a TissueLyser LT (Qiagen) in RIPA buffer (Sigma R0278). Protein concentration was determined by Biorad Protein Assay (Biorad) (Biorad), lysate protein concentration adjusted to 0.5 µg/µl protein liver lysate and submitted to the Human Immune Monitoring Center (HIMC) a core facility at Icahn School of Medicine at Mount Sinai.

Olink data were generated using the Olink mouse exploratory panel (v.3801), and the data was exported in NPX format from the Olink NPX Manager (v. 2.0.0.173) to Excel xlsx files. These files were imported into an R session (development version, 2022-01-07 r81454) using the OlinkAnalyze package (v. 3.0.0). Groups were compared using a one-way ANOVA with Tukey’s post-hoc test as implemented in the OlinkAnalyze package and using the default adjusted P value of 0.05. Other R packages used to generate figures and plots were the packages NMF (v. 0.23.0), ggplot2 (v. 3.3.5), and cowplot (v. 1.1.1).

#### Immunohistochemistry

For immunohistochemistry, formalin-fixed paraffin embedded sections were cut at 4 µm thickness and slides incubated at 65°C for 30 min. Slides were deparaffinized in Histoclear (National Diagnostics, HS2001GLL) and rehydrated by an ethanol gradient. Heat antigen retrieval was performed in 10 mM sodium citrate (pH 6.0) in a pressure cooker (Cuisinart, CPC-600C), 20 min at high-pressure, natural pressure release for 30 min, and cooled at room temperature for 30 min. Endogenous peroxidase quenching was performed with 1.5% hydrogen peroxide (Sigma-Aldrich, H1009) for 20 min, and slides washed twice in phosphate buffered saline (PBS) twice. Blocking was performed for 30 min with Protein Block, Serum-free (Agilent, X090930-2). The primary antibody (**Table S2**) was diluted in antibody diluent (Agilent, S080983-2) and applied to the slides in a wet chamber overnight at 4°C. After 2 washes in PBS, slides were incubated with the appropriate secondary antibody (**Table S2**) for 30 min followed by two washes in PBS. The antigen was revealed with DAB (Vector Laboratories, SK-4105) until the appropriate staining on a control slide was obtained. The same reveal time was then used on all experimental slides. Slides were stained with hematoxylin (Sigma-Aldrich, GHS332) for 30 sec, washed gently in lukewarm tap water for 5 min, and washed twice in distilled water, followed by two washes in phosphate buffered saline (PBS). Slides were washed in increasing ethanol/water gradients, followed by three washes in Histoclear. Whole-slide scans for subsequent analyses and publishing were obtained using an VS200 Research Slide Scanner (Olympus, Tokyo, Japan) and a SCN400 (Leica, Wetzlar, Germany). Whole-slide images were processed and annotated in QuPath 0.3.2. Ki67-positive hepatocyte nuclei were counted over ten random high-power fields (ROIs of 0.27 mm^2^). Hepatocytes and cleaved caspase 3-positive hepatocyte nuclei were counted over 5 random high-power fields and mean % of cleaved caspase 3-positive hepatocytes per mouse calculated.

#### Immunofluorescence

For immunofluorescence, 4 µm cryosections from fresh frozen liver samples embedded in Optimal Cutting Temperature (OCT) compound (Sakura, Tokyo, Japan) were cut and dried for a minimum of 1 h at room temperature and stored at -80°C until use. Frozen sections were thawed and rehydrated in PBS for 5 min at room temperature, fixed in 100% acetone for 10 min at -20°C and washed in PBS. The primary antibody (**Table S2**) was diluted in antibody diluent (Agilent, S080983-2) and incubated for 60 min at room temperature in a wet chamber. Slides were then washed 3 x 5 min in PBS. The secondary antibody (**Table S2**) was diluted in antibody diluent (Agilent, S080983-2), and slides incubated in a wet chamber for 30 min. Slides were washed three times in PBS, followed by two washes in distilled H_2_O. The slides were then mounted with a DAPI-containing medium (Prolong Antifade, Thermo Fisher Scientific, P36931). Images were taken with either a Nikon A1R confocal microscope (Nikon, Tokyo, Japan), or a Axioimager M2, or a Axioimager 7 (Zeiss, Oberkochen, Germany) microscope.

#### Multiplex immunohistochemistry

For multiplex immunohistochemistry, FFPE sections were incubated at 65°C for 30 min and cooled for 15 min at room temperature for 2 cycles to strongly fix the tissue to the slide. After deparaffinization and rehydration, heat antigen retrieval was performed with 10 mM sodium citrate (pH 6). The slides were then stained with hematoxylin to avoid staining after the signal reveal which would unbind the chromogen. Endogenous peroxidase quenching was performed with 1.5% hydrogen peroxide (Sigma-Aldrich, H1009) for 20 min, and slides washed twice in phosphate buffered saline (PBS). Blocking was performed for 30 min with Protein Block, Serum-free (Agilent, X090930-2). The CCND1 primary antibody (**Table S2**) was diluted in antibody diluent (Agilent, S080983-2) and incubated on the slides for 60 min. Slides were washed in PBS and incubated with secondary antibody for 30 min. Slides were washed in PBS and slides incubated with 3-amino-9-ethylcarbazole (AEC, Agilent, K346111-2) until the appropriate staining on a control slide was obtained. The same reveal time was used on all experimental slides. The slides were washed in PBS, and cover slipped with a PBS mounting medium immediately prior to imaging. Slides were scanned with the VS200 research slide scanner (Olympus, Tokyo, Japan). After imaging, the cover glass were carefully removed in a PBS wash, and the slides were de-stained in a gradient of dH_2_O, 70% ethanol, 95% ethanol, 70% ethanol, dH_2_O, and conserved in PBS at 4°C. Slides were incubated in 10 mM sodium citrate (pH 6.0) in a pressure cooker (Cuisinart, CPC-600C), for 20 min at high-pressure, natural pressure release for 30 min, and cooled at room temperature for 30 min. The hematoxylin staining was repeated, and the steps were repeated from the blocking step but with anti-HNF4α antibody (**Table S2**). Images were registered in QuPath 0.3.2 with the image combiner extension. Ten high-power fields (ROIs measuring 0.27 mm^2^) evenly distributed across tissue sections per sample were assessed.

#### Cell lines

Human HepG2 (ATCC HB-8065) were maintained in MEM media with 10% FBS and the mouse hepatocyte cell line AML12 (ATCC CRL-2254) in DMEM/F12 media supplemented with dexamethasone (40 ng/ml) and Insulin-Transferrin-Selenium (Gibco, 41400045), containing 10% FBS and 1% penicillin and streptomycin. JS1, LX2 and TWNT4 cells were cultured in DMEM supplemented with 10% FBS.

#### Primary hepatocyte isolation

*Pdgfrb*-GFP-negative or H-2K^d^ mice older than 4 months were anesthetized and cannulated via the portal vein. Livers were perfused with ∼25 ml of pre-warmed Liver Perfusion Medium (Gibco, 17707-038) followed by ∼50 ml Liver Digestion Medium (Gibco,17703-034). Liver capsule was gently teased apart with tweezers and hepatocytes released. Cells were taken up in Hepatocyte Wash Buffer (Gibco, 17704-024) and filtered through 100 µm mesh. Cells were then centrifuged for 2 min at 100g, 4°C and cell pellet resuspended in ice cold 10 ml DMEM (Gibco, 11885-084)/5% FBS. 10 ml of ice cold 100% Percoll/PBS (GE, GE17-0891-01) was added for gradient purification and cells centrifuged for 10 min at 200g, 4°C. Cell pellet was resuspended with 20 ml DMEM/5%FBS and centrifuged for 2 min at 100g, 4°C. 1-2 x 10^3^ hepatocytes were plated on collagen-coated 96-well plates in DMEM/5%FBS/Pen/Strep. Rat tail collagen I (Gibco, A1048301) was used for coating. Cell culture media was changed 3 hours after plating with either serum-free William’s E media (Gibco) containing Pen/Strep (Gibco) or DMEM/5%FBS/Pen/Strep. NTF3 (STEMCELL technologies) was added 1 day after plating at a final concentration of 20 ng/ml and primary hepatocytes cultured for 3 days with NTF3 until analysis.

#### Cell proliferation assays

Primary mouse hepatocytes, HepG2 or AML12 cells were grown in opaque-walled 96-well plates. NTF3 (final concentration 20 ng/ml) (STEMCELL Technologies, cat. 78074) and/or Loxo-195 (final concentration 5 nM) (Selleckchem, S8636) were added into complete growth media without L-glutamine. Cells were analyzed by CellTiter Glo (Promega G7570, Madison, WI) which detects the number of viable cells on day 0 to day 5 by on detection systems Glomax (Promega) or Bioteck Synergy h1 (Agilent).

#### Western blotting

Fifty micrograms of snap frozen liver were homogenized with a Qiagen PowerLyser in RIPA buffer containing proteinase inhibitors (Roche complete Mini proteinase inhibitor, 2 tab/10 ml buffer) and 1x HALT phosphatase inhibitors (Thermoscientific, cat. 78428). Lysates were sonicated and then centrifuged for 10 min at 18,800 × g. Protein concentration of supernatants were determined by Direct Detect Spectrophotometer (Millipore).

Human HepG2 and mouse AML12 cells were grown in 6-well plates. Cells were treated with NTF3 (20 ng/ml) purchased from STEMCELL technologies (cat. 78074), R&D (267-N3), Gibco (PHC7036) and/or Loxo-195 (5-10 nM) (Selleckchem, S8636) in complete growth media without L-glutamine for 2 hours. Lyophilized NTF3 was solubilized with sterile filtered PBS supplemented with 0.1% BSA, aliquoted and stored at -80°C until use. Only aliquots with < 3 months shelf life were used for experiments as activity and protein concentration declined significantly after 3 months. HepG2 and AML12 cells were collected on ice in PBS, centrifuged for 10 sec at and cell pellet homogenized in RIPA buffer containing inhibitors (Roche complete Mini proteinase inhibitor, 1x HALT phosphatase inhibitors, Thermoscientific, cat. 78428). Cell lysates were sonicated and centrifuged for 10 min at 18,800 × g. Protein concentration of supernatants were determined by Direct Detect Spectrophotometer (Millipore).

For immunoblot analysis, 30–50 μg of protein diluted in LSD sample buffer (Invitrogen) were analyzed per SDS-PAGE (Nupage, Invitrogen) using MOPS buffer or MES buffer (Invitrogen). Gels were stained with Coomassie (Bio-Safe Coomassie G-250 Stain, Biorad) and transferred onto PVDF membranes using semi-dry blotting (Trans-Blot SD, Biorad). Antibodies and dilutions used are indicated in **Table S3**. Images were visualized on an Amersham Imager 680 (Cytiva). Semi-quantitative band densitometry was performed using ImageJ software (version 1.53).

#### Statistical Analysis

For analysis of the Olink and RNA-seq data see the corresponding sections. Other data analysis was performed using GraphPad Prism (version 9.4.1). Two groups were compared by unpaired two-tailed Student’s t-test. Data of three or more groups were first assessed for normality and then either assessed by one-way ANOVA with the post hoc tests by Tukey, Kruskal-Wallis, or Dunn.

---

**Suppl. Table 1.** Gene set analysis of metabolic pathways from HSC-depleted livers and controls. Self-contained gene set pathway analysis for gene sets and pathways defined in Reactome, Biocarta, and KEGG databases comparing bulk liver RNA-seq of HSC-depleted and control mouse livers 10 days after adoptive transfer (n=4 vs. n=6 respectively). The Bioconductor edgeR::mroast function (version 3.37.0) (*60*) was used to test the mouse MSigDB C2 gene set collection. It was downloaded from https://bioinf.wehi.edu.au/software/MSigDB/.

**Supplemental Table S2.** List of primary and secondary antibodies for immunohistochemistry, immunofluorescence, and multiplex immunohistochemistry.

| Name | Cat# | Vendor | Application (IHC/IF) | Dilution |
| --- | --- | --- | --- | --- |
| cleaved caspase 3 | 9661 | Cell signaling | IHC | 1:3000 |
| CCND1 | ab16663 | Abcam | IHC | 1:800 |
| CD31/Pecam1 | 550274 | BD Pharmingen | IHC | 1:100 |
| desmin (rabbit host) | ab15200 | Abcam | IHC | 1:8000 |
| desmin (goat host) | AF3844 | R&D | IF | 1:25 |
| E-cadherin | 3195S | Cell Signaling | IHC | 1:3000 |
| F4/80 | 14-4801-82 | eBioscience | IHC | 1:100 |
| GFAP | NB300-141 | Novus Biologicals | IHC<br>IF | 1:1000<br>1:50 |
| GFP | ab183735 | Abcam | IHC<br>IF | 1:1000<br>1:200 |
| glutamine synthetase | ab16802 | Abcam | IHC | 1:200 |
| granzyme B | ab4059 | Abcam | IHC | 1:5000 |
| HNF4α | ab181604 | Abcam | IHC | 1:400 |
| Ki67 | M7249 | Dako | IHC | 1:50 |
| NTF3 | PA5-14861 | Invitrogen | IHC<br>IF | 1:500<br>1:50 |
| TRKA (*NTRK1*) | AF1056SP | R&D | IF | 1:100 |
| TRKB (*NTRK2*) | AF1494 | R&D | IF | 1:100 |
| reelin | AF3820 | R&D | IF | 1:50 |
| Anti-Mouse IgG, Secondary antibody, Peroxidase | MP-7402 | Vector Laboratories | IHC | 1:2000 |
| Goat Anti-Rat IgG, Secondary antibody, Peroxidase | MP-7404 | Vector Laboratories | IHC | 1:2000 |
| Goat anti-Rabbit IgG,<br>Secondary antibody,<br>Peroxidase | MP-7451 | Vector<br>Laboratories | IHC | 1:2000 |
| Donkey Anti-Goat<br>Secondary Antibody, Cy3 | A-21447 | Invitrogen | IF | 1:2000 |
| Goat anti-Rabbit Secondary<br>Antibody, Alexa Fluor 647 | A-21245 | Invitrogen | IF | 1:2000 |

**Supplemental Table S3.** List of primary antibodies for western blotting.

| **Name** | **Cat#** | **Vendor** | **Dilution** |
| --- | --- | --- | --- |
| β-actin | ab6276 | Abcam | 1:10,000 |
| CCND1 | ab16663 | Abcam | 1:750 |
| desmin | ab15200 | Abcam | 1:1000 |
| IGFBP2 | ab188200 | Abcam | 1:1000 |
| tubulin | 3873 | Cell signaling | 1:10,000 |
| GAPDH | CB1001 | Calbiochem | 1:5,000 |
| NTF3 | PA1-18385 | Invitrogen | 1:1000 |
| PCNA | 13110 | Cell signaling | 1:1000 |
| retinoblastoma | 9309 | Cell signaling | 1:1000 |
| Phospho-retinoblastoma | 8180 | Cell signaling | 1:1000 |
| Pan-TRK | 92991S | Cell signaling | 1:1000 |
| Phospho-TRK (P-TRKA<br>(Tyr674/675)/TrkB (Tyr706/707)) | 4621S | Cell signaling | 1:1000 |

